# A multisensory circuit for gating intense aversive experiences

**DOI:** 10.1101/2021.05.01.441648

**Authors:** Arun Asok, Félix Leroy, Cameron Parro, Christopher A. de Solis, Lenzie Ford, Michelle N. Fitzpatrick, Abigail Kalmbach, Rachael Neve, Joseph B Rayman, Eric R. Kandel

## Abstract

The ventral hippocampus (vHPC) is critical for both learned and innate fear, but how discrete projections control different types of fear is poorly understood. Here, we report a novel excitatory circuit from a subpopulation of the ventral hippocampus CA1 subfield (vCA1) to the inhibitory peri-paraventricular nucleus of the hypothalamus (pPVN) which then routes to the periaqueductal grey (PAG). We find that vCA1→pPVN projections modulate *both* learned and innate fear. Fiber photometric calcium recordings reveal that activity in vCA1→pPVN projections increases during the first moments of exposure to an unconditioned threat. Chemogenetic or optogenetic silencing of vCA1→pPVN cell bodies or vCA1→pPVN axon terminals in the pPVN enhances the initial magnitude of both active and passive unconditioned defensive responses, irrespective of the sensory modalities engaged by a particular innate threat. Moreover, silencing produces a dramatic impact on learned fear without affecting milder anxiety-like behaviors. We also show that vCA1→pPVN projections monosynaptically route to the PAG, a key brain region that orchestrates the fear response. Surprisingly, optogenetic silencing of vCA1 terminals in the pPVN titrates the level of c-Fos neural activity in the PAG differently for learned versus innate threats. Together, our results show how a novel vCA1→pPVN circuit modulates neuronal activity in the PAG to regulate both learned and innate fear. These findings have implications for how initial trauma processing may influence maladaptive defensive behaviors across fear and trauma-related disorders.

**One Sentence Summary:** A multisensory gate for high intensity aversive experiences.

## Introduction

Fear- and trauma-related disorders affect tens of millions of people each year (*1*) – a statistic likely amplified by the COVID pandemic (*2*). How we initially respond to, and later recall, a traumatic experience can exert a powerful influence over our future behavior (*3-5*). Yet, our current understanding of the neural circuits that control learned and innate fears is incomplete (*6-8*). It has long been appreciated that the amygdala plays a central role in the regulation of fear memory, with many studies having elucidated which amygdalar cell-types, subdivisions, and circuits control learned versus innate fear (for review see (*9, 10*)). By contrast, we are just beginning to appreciate how the ventral hippocampus also controls different types of fear and anxiety-related behaviors (*6, 11-13*), as well as how the coordinated interplay between neural circuits across different brain regions gives rise to adaptive or maladaptive behavior (*14*).

While the function of the ventral hippocampus in learned fear is partly characterized (for review see (*11*)), its role in innate fear and anxiety is less clear (*11, 15*). For example, electrolytic and neurotoxic lesions of the ventral hippocampus disrupt freezing to conditioned cues and contexts as well as inhibitory avoidance (*16-18*). Importantly, ventral hippocampal lesions also produce a marked impairment in anxiety (*15*) as well as an impairment in fear to predatory odors, but the precise function of the ventral hippocampus in innate predatory fear is poorly understood (*19*). The diverse role of the ventral hippocampus in different types of fear and anxiety-related behaviors highlights a more global function across various defensive behaviors. That is, the ventral hippocampus may control defensive behaviors by modulating core innate or unconditioned aspects of an aversive experience (*15, 20*).

The function of the ventral hippocampus in guiding defensive behaviors is mediated by discrete outputs. For instance, reciprocal circuits between the ventral hippocampus and the basolateral amygdala control conditioned fear and anxiety-related avoidance to a context (*12, 21-23*) whereas ventral hippocampal projections to the central amygdala are important for learned fear to discrete cues (*12*). Moreover, ventral hippocampal projections to the medial prefrontal cortex and the lateral hypothalamus control anxiety (*13, 22, 24*). Despite tremendous progress in identifying the function of specific ventral hippocampal circuits, it is still unclear how the ventral hippocampus controls both innate and learned defensive behaviors.

Here, we use a combination of viral circuit targeting, whole-cell patch clamp, chemogenetics, fiber photometry, optogenetics, and various cell-type specific labeling methods to elucidate how a novel ventral hippocampal to hypothalamic projection controls learned and innate fear. Our findings reveal fundamental insights into how the ventral hippocampus processes learned and innate aversive information to influence defensive responses to a variety of threats.

## Results

We hypothesized that the ventral hippocampus regulates learned and innate fear through projections to the hypothalamus, a brain region whose anterior, ventromedial, and paraventricular sub-division are known to regulate different aspects of learned fear, innate fear, and homeostatic neuroendocrine processes (*25-29*). To reveal this circuitry, we injected an adeno-associated virus (AAV) under control of the calcium-calmodulin dependent protein kinase II α promoter expressing eYFP (AAV8-CAMKIIα-eYFP) into the ventral hippocampus to determine where long-range excitatory projections to the hypothalamus cluster (Fig. 1a). Viral tracing revealed eYFP^+^ fibers surrounding the paraventricular nucleus of the hypothalamus (PVN), where vCA1 projections formed a halo-like pattern in the peri-paraventricular area (pPVN) that surrounds the dense 60-100um thick neuronal population that characterizes the PVN (Fig. 1b-c; (*30*). Retrograde G-deleted rabies virus tracing from the PVN revealed that vCA1 projections to the pPVN are monosynaptic (fig. S1a-c). Given the halo-like distribution of pPVN neurons around the PVN, we refer to pPVN^GAD67^ neurons as Halo Cells.

**Figure 1.**
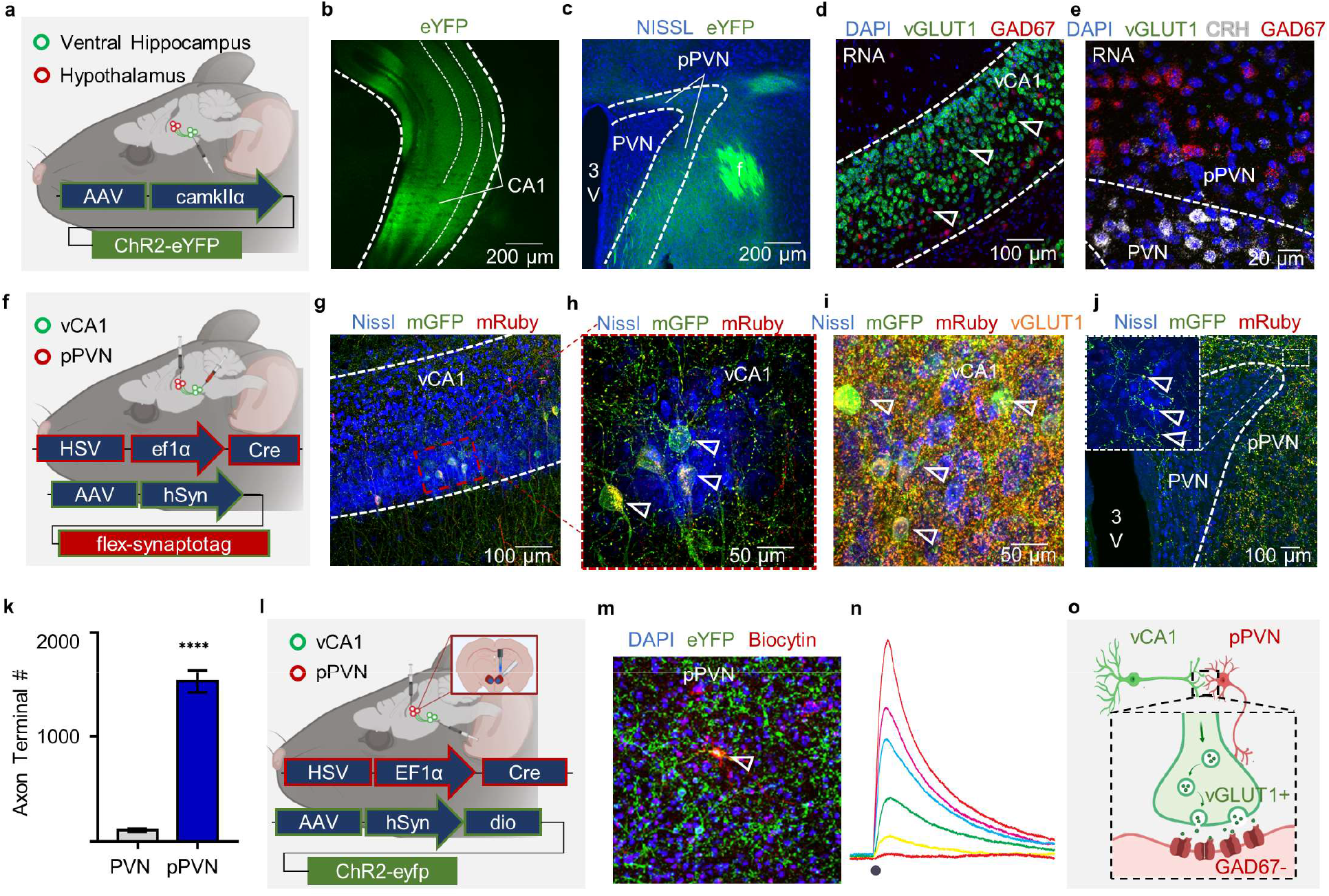
The ventral CA1 subfield of the hippocampus (vCA1) projects to the peri-paraventricular nucleus of the hypothalamus (pPVN). **(a)** schematic of AAV-CAMKII-ChR2-EYFP single-site viral injection into vCA1. **(b)** ChR2-eYFP (green) expression in ventral hippocampus. **(c)** ChR2-eYFP (green) expression in the pPVN. Note the “halo-like” pattern around the dense Nissl stained (blue) neurons in the paraventricular nucleus of the hypothalamus (PVN). **(d)** in situ hybridization identifying vCA1 cells (blue) predominantly express vGlut1 (green) and little GAD67 (red). **(e)** in situ hybridization of PVN area identifying that the corticotropin-releasing hormone (CRH; grey) expressing cells (blue) are surrounded by GAD67 (red). Note the lack of vGlut1 labeling (green). **(f)** schematic of dual-site viral injection of retrograde HSV-EF1α-Cre into the pPVN followed by a Cre-dependent AAV expressing mGFP and synaptophysin fused to mRuby (i.e., AAV-synaptotag). **(g)** expression of mGFP (green) and synaptophysin-mRuby (red) in neurons (blue) in vCA1. **(h)** magnified image showing mGFP (green) and mRuby (red) expression throughout vCA1→pPVN projecting neurons. **(i)** mGFP^+^ (green) and mRuby (red) vCA1→pPVN projecting neurons (blue) express vGlut1 (orange). (j) vCA1→pPVN axonal projections (green) predominantly synapse in the pPVN (red) relative to the dense cluster of PVN neurons (blue). Inset shows magnified view of terminals in the pPVN. **(k)** vCA1 projecting neurons exhibit significantly more synapses in the pPVN (blue bar) relative to the PVN (grey bar; *t*_4_ = 22.64, p < 0.0001, 12 slices from 3 mice). **(l)** schematic of optogenetic whole-cell patch clamp approach following injections of HSV-Cre-mCherry into the pPVN followed by injection of a Cre-dependent AAV-ChR2-eYFP construct in vCA1. **(m)** Biocytin-filled neuron (red) surrounded by ChR2-eYFP expressing fibers (green) in the pPVN. **(n)** voltage response from of pPVN neuron during photoactivation of ChR2-expressing vHPC inputs using increasing light intensity. Inset shows response of same pPVN cell during injection of current pulses. **(o)** Schematic of excitatory vGlut1^+^ vCA1 neurons projecting to inhibitory GAD67^+^ neurons in the pPVN. Bar graphs are mean ± S.E.M.

To identify the excitatory and inhibitory organization of vCA1, PVN, and pPVN neurons, we performed dual- and triple-labeled fluorescent *in situ* hybridization. Neurons in vCA1 were primarily vGlut1-positive (vGlut1^+^; Fig. 1d) and PVN neurons were corticotropin-releasing hormone (CRH; a neuropeptide selectively enriched in the PVN (*31*)) positive but not vGlut1^+^, whereas pPVN neurons were GAD67^+^, but not CRH^+^ or vGlut1^+^ (Fig. 1e; (*32*)). In order to confirm that vCA1 projections synapsed in the pPVN, we injected a monosynaptic retrograde herpes-simplex virus harboring Cre-recombinase (HSV-Cre) into the PVN followed by a Cre-dependent adeno-associated virus (AAV) expressing a membrane-bound variant of GFP and synaptophysin-mRuby to fluorescently label axon terminals in the pPVN from vCA1 neurons (Fig. 1f). This dual viral approach allowed for selective targeting of the vHPC→pPVN projections. Immunohistochemical labeling confirmed that vCA1→pPVN neurons were in fact vGlut1^+^, while mRuby^+^ axon terminals localized primarily in the pPVN relative to the PVN (1530.7 ± 62.2 versus 107.3 ± 9.6 terminals; Fig. 1g-k). Taken together, these results confirm that excitatory vCA1 neurons project to inhibitory neurons in the pPVN.

vCA1 neurons are known to project to both the amygdala and medial prefrontal cortex, and both of these projections regulate fear and anxiety (*12, 22*). Thus, we sought to determine if vCA1→pPVN projections were collaterals of bifurcating vCA1 projection neurons. We used a dual retrograde HSV approach to examine the overlap between vCA1→pPVN projections (318 neurons, 16 mice) and other known ventral hippocampal projections including: vCA1→basolateral amygdala (153 neurons, 8.46% overlap, 4 mice), vCA1→medial prefrontal cortex (1103 neurons total, 0.47% overlap, 4 mice), vCA1→lateral hypothalamus (466 neurons, 0.42% overlap, 4 mice), and vCA1→nucleus accumbens (522 neurons, 0.64%, 4 mice). Immunohistochemical analysis revealed little to no overlap with other known vCA1 circuits, suggesting vCA1 neurons projecting to the pPVN form a non-branching distinct neuronal population (Fig. S1 d-o). Taken together, our results suggest that a subpopulation of vCA1^vGlut1^ neurons send monosynaptic inputs to inhibitory pPVN^GAD67^ neurons.

Next, to dissect how vCA1→pPVN projections influence postsynaptic pPVN targets, we injected retrograde HSV-Cre in the PVN area followed by a Cre-dependent AAV expressing channelrhodopsin fused with eYFP into vCA1 (AAV8-hSYN-DIO-ChR2-eYFP). We prepared acute slices of the PVN area and obtained whole-cell patch clamp recordings of pPVN cells (Fig. 1m-n & S2a). Light-stimulation induced strong mono-synaptic EPSPs (Fig. 1n) that could trigger action potentials (not shown). Bath application of glutamatergic synaptic blockers abolished the response whereas application of GABAergic synaptic blockers potentiated it (Fig. s2), suggesting the existence of feed-forward inhibition between pPVN neurons. Moreover, the short latency of the light-induced EPSPs in post-synaptic targets further confirmed the monosynaptic nature of the vCA1→pPVN pathway (Fig. 1n). By contrast, PVN cell recordings showed no light-induced response (not shown). Overall, these data show that vCA1 neurons projecting to the PVN area excite pPVN inhibitory neurons (Fig. 1o).

How do vCA1→pPVN projections control defensive behavior? Given that the ventral hippocampus is known to regulate conditioned fear (*15*), unconditioned predator odor fear (*19*), and anxiety-like behaviors (*22*), we hypothesized that the vCA1→pPVN projections may play a critical role in modulating a core component (i.e., unconditioned stimulus (US) processing) between these different experiences. To test this, we used a chemogenetic approach and combined injections of retrograde HSV-Cre into the pPVN with injections of a Cre-dependent AAV8 harboring hm4di fused to mCherry (hm4di) or mCherry-only (mCherry) as a control into vCA1 (Fig. 2a & S3a). We administered the hm4di agonist clozapine-N-oxide (CNO) 20-m prior to behavioral testing. Chemogenetic silencing had no effect on anxiety-like behaviors in the open field (Fig. S3b-f) or elevated plus maze tests (EPM, Fig. S3g-k). Moreover, chemogenetic silencing did not affect locomotor activity in the open-field test (Fig. S3c-d). Finally, although silencing had no effect on acoustic startle responsivity or short-term startle habituation (Fig. S3l-o), silencing preferentially disrupted pre-pulse inhibition to the highest pre-pulse stimulus (Fig. S3n). Taken together, these data suggest that while the vCA1→pPVN circuit is not important for regulating milder anxiety-like behaviors, it is important for behaviorally gating the defensive response to more intense aversive experiences.

**Figure 2.**
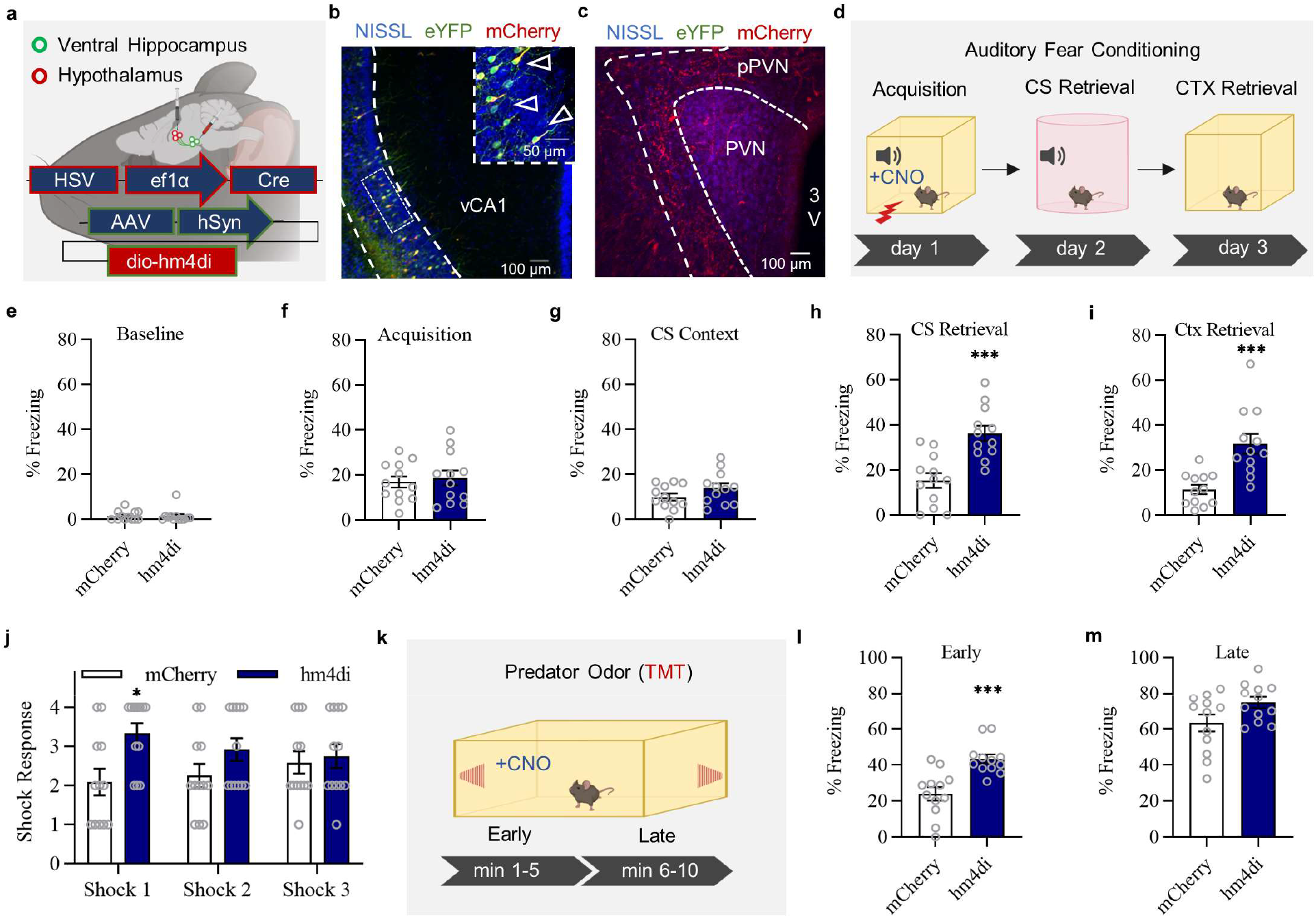
Chemogenetic silencing of vCA1→pPVN projections enhances learned and innate fear by preferentially modulating reactivity to unconditioned threats. **(a)** schematic of dual-site viral injection approach using a retrograde HSV-EF1α-Cre-eYFP injected into the pPVN followed by a Cre-dependent AAV expressing hM4Di-mCherry or mCherry alone injected into vCA1. **(b)** representative image of co-localization between HSV-EF1α-Cre-eYFP and AAV-hSyn-DIO-hM4Di-mCherry in vCA1. Inset is magnified view of co-localization from panel B. **(c)** representative image of vCA1 mCherry^+^ axons in the pPVN. **(d)** Schematic of 3-day auditory fear conditioning paradigm. Mice were administered Clozapine-N-oxide (CNO) 20 min prior to conditioning. **(e)** CNO had no effect in any group on baseline freezing to the conditioning context: (t_22_=0.110, p= 0.9126), **(f)** freezing during acquisition (t_22_=0.4393, p= 0.6647), or **(g)** baseline freezing in the novel CS testing context (t_22_=1.536, p= 0.1388). **(h)** hM4Di + CNO mice exhibited enhanced cue-specific freezing during non-reinforced exposure to conditioned CSs in a novel context, one-way (t_22_=4.467, p= 0.0002). (i) hm4di + CNO mice showed elevated freezing during non-reinforced re-exposure to the original conditioning context, (t_22_=4.212, p= 0.0004). **(j)** hM4Di + CNO mice exhibited enhanced shock responsivity to the first foot-shock during conditioning (Mann-Whitney_2-tailed_ U = 29.5, p = 0.0105), but not subsequent shocks (p > 0.05). **(k)** schematic of predator odor exposure paradigm highlighting early (1-5 min) and late phase (5-10 min) of the test. **(l)** hM4Di + CNO mice exhibited enhanced freezing during the first five minutes of exposure to the predator odor 2,5-dihydro-2,4,5-trimethylthiazoline (TMT), (t_22_=4.317, p= 0.0003). **(m)** hM4Di + CNO mice did not differ from other groups during the last five minutes of exposure to TMT, (t_22_=2.032, p= 0.0544). Bars and bar graph error bars are mean ± S.E.M. *p < 0.05, ***p < 0.001. conditioning: n = 12 mice/group, odors: n = 12 mice/group.

Therefore, based on these results, we hypothesized that chemogenetic silencing of vCA1→pPVN projections may preferentially influence higher-intensity unconditioned aversive experiences. Thus, we used the same chemogenetic strategy to silence vCA1→pPVN projections prior to auditory delay fear conditioning (Fig. 2a-d) and found no effect on locomotor activity/baseline freezing at conditioning, acquisition, or future baseline freezing in the conditioned stimulus (CS) testing context during cue-retrieval (Fig. 2e-g). However, silencing enhanced CS-specific freezing in a novel context and background contextual freezing in the original conditioning context (Fig. 2h-i). Given that silencing (1) only influenced startle at the highest pre-pulse during pre-pulse inhibition, (2) enhanced freezing during auditory fear retrieval, but (3) did not affect performance-related freezing at baseline or acquisition, we reasoned that the vCA1→pPVN circuit may modulate auditory fear memories by affecting unconditioned foot-shock responses. Surprisingly, silencing vCA1→pPVN projections preferentially enhanced responsivity to the first foot-shock, but not subsequent foot-shocks (Fig. 2k).

Given that (1) the core sensory input pathways differ between pre-pulse inhibition and foot-shock (i.e., auditory vs. tactile), in addition to (2) constraints across intensity (i.e., the highest pre-pulse) and (3) time (i.e., first foot-shock during fear conditioning) domains, we reasoned that vCA1→pPVN projections may gate unconditioned defensive responses to more intense aversive experiences in a temporally dependent manner. That is, activity would scale during the initial seconds of the first foot-shock during conditioning and the first minutes during sustained presentation of a US through other sensory modalities. To test this hypothesis, we leveraged a predator odor exposure paradigm to probe the effect of chemogenetic silencing on unconditioned olfactory fear (Fig. 2l). vCA1→pPVN silencing enhanced unconditioned freezing to the predator odor 2,5-dihydro-2,4,5-trimethylthiazoline (TMT) during early (minutes 1-5; Fig. 2m), but not later (minutes 5-10) time periods (Fig. 2n). Chemogenetic silencing had no effect on overall performance given that mice were able to reach asymptotic levels of freezing during the later parts of the exposure session (Fig. 2n). Taken together, these data suggest that vCA1→pPVN projections temporally gate defensive freezing to an unconditioned olfactory threat. Moreover, these findings extend upon the previous fear conditioning and PPI experiments to indicate that the vCA1-pPVN circuit preferentially controls higher-intensity unconditioned aversive experiences, irrespective of the sensory-modality through which unconditioned/innate aversive information enters the brain.

Does vCA1→pPVN calcium activity parallel our behavioral findings and increase during the first foot-shock of auditory-fear conditioning or the early-parts of exposure to a predator odor? To answer this, we injected a retrograde HSV construct expressing the genetically encoded calcium indicator GCaMP6f into the pPVN and implanted an optical fiber above vCA1 to record calcium activity from vCA1→pPVN projecting neurons during auditory fear conditioning (Fig. 3a-c). In accordance with our behavioral data, vCA1 neurons projecting to the pPVN exhibited a significant increase in calcium activity during the first, but not during subsequent, foot-shocks. Calcium responses were highly elevated during the first shock with the only significant increases detected during the first few minutes (60 to 120s) of long-term fear memory tests (Fig. 3e-g). These data suggest that the vCA1→pPVN circuit may have a modulatory role in gating innate/unconditioned fear (given temporally restricted calcium activity during non-reinforced exposure to the CS in a novel context or the background conditioning context). Interestingly, exposure to the novel CS testing context or original conditioning context produced a sustained elevation in calcium activity during the first minutes of exposure – providing support for recent computational models on how the hippocampus functions (*33, 34*). Finally, exposure to the predator odor TMT resulted in a sustained elevation of calcium activity during the early (first 150s), but not later phase of exposure, similar to our behavioral results (Fig. 3h-j). The chemogenetic data along with our *in vivo* calcium recordings suggest that the vCA1→pPVN circuit contributes to temporally gating the initial defensive response to different types of unconditioned threats. Furthermore, vCA1→pPVN may participate during the stimulus sampling phase of sensory contextual information, irrespective of whether the context is aversive.

**Figure 3.**
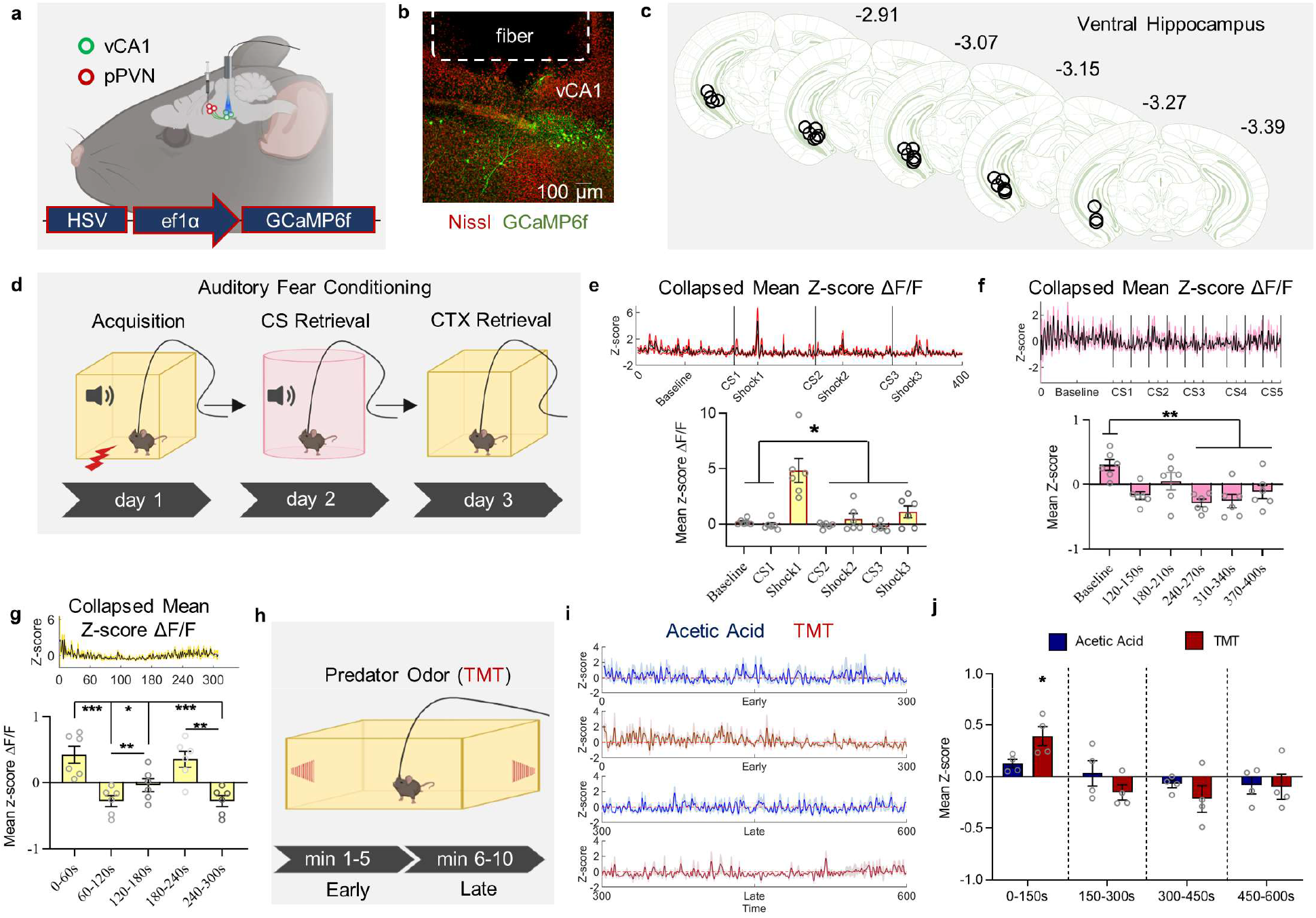
Fiber photometric calcium recordings of vCA1→pPVN projections exhibit temporally-restricted activity to unconditioned/innate stimuli. **(a)** schematic of pPVN viral injection HSV-EF1α-GCaMP6f and optical fiber implant in vCA1. **(b)** representative image of vCA1 fiber optic tract and GCaMP6f expression in vCA1. **(c)** reconstruction of fiber optic placements. (d) schematic of auditory fear conditioning paradigm. **(e)** top panel: collapsed calcium activity across all mice during acquisition. bottom panel: mean z-score transformed ΔF/F (ANOVA_1-way Brown-Forsythe corrected_: F_6,35_ = 3.484, p = 0.0083). **(f)** top panel: collapsed calcium activity across all mice during non-reinforced CS testing in a novel context, bottom panel: mean z-score transformed ΔF/F where baseline activity was significantly elevated over all other phases (ANOVA_1-way_: F_5,30_ = 5.483, p = 0.0011). **(g)** top panel: collapsed calcium activity across all mice during non-reinforced testing in the conditioning context, bottom panel: mean z-score ΔF/F where baseline activity was elevated during the early parts of testing (ANOVA_1-way_: F_4,25_= 10.26, p < 0.0001). **(h)** schematic of predator odor fear paradigm. **(i)** collapsed calcium activity during early (top panels) and late (bottom panels) parts of odor exposure to acetic acid or TMT. **(j)** mean z-score ΔF/F where calcium activity was elevated during the first 150-s of exposure (ANOVA_2-way main effect of group_: F_3,12_ = 19.61, p<0.0001). Shaded error bars and bar graph error bars are mean ± S.E.M. conditioning: n = 6 mice, odors: n = 4 mice/group.

We next sought to examine how vCA1→pPVN projections could gate defensive responses by identifying which brain area the pPVN primarily communicates with. To this end, we designed an AAV expressing Cre/eYFP under the control of a DLX gene promoter specific to inhibitory neurons (AAV9-DLX-Cre-eYFP), with the goal of targeting inhibitory neurons in the pPVN (Fig. S4a). Using immunohistochemistry (IHC) and *in situ* hybridization (ISH), we confirmed that our viral strategy recapitulates the pPVN expression pattern observed by vCA1 projections cells (Fig. S4b) and specifically targets GAD67^+^ pPVN cells (Fig. S4c-d). Surprisingly, pPVN neurons did not exhibit clear projections to the PVN (Fig. S4b) – providing evidence against the long-held hypothesis that pPVN neurons provide direct feed-forward inhibition of the PVN (*35, 36*). If not local, we hypothesized that the pPVN may therefore control passive and active defensive behaviors through long-range inhibitory inputs to other regions such as the periaqueductal grey (PAG; (*37, 38*). Indeed, when we co-injected AAV9-DLX-Cre-eYFP and AAVDJ-Flex-synaptotag into the pPVN we found a robust pPVN projection to the dorso-lateral portion of the PAG (dlPAG; Fig. 4a-b). Given that dorsolateral (dl) and ventrolateral (vl) divisions of the PAG are critical for active and passive defensive responses (*39*), respectively, these data suggested that the vCA1→pPVN→PAG circuit may provide a route by which vCA1→pPVN silencing influences both learned and innate fear.

**Figure 4.**
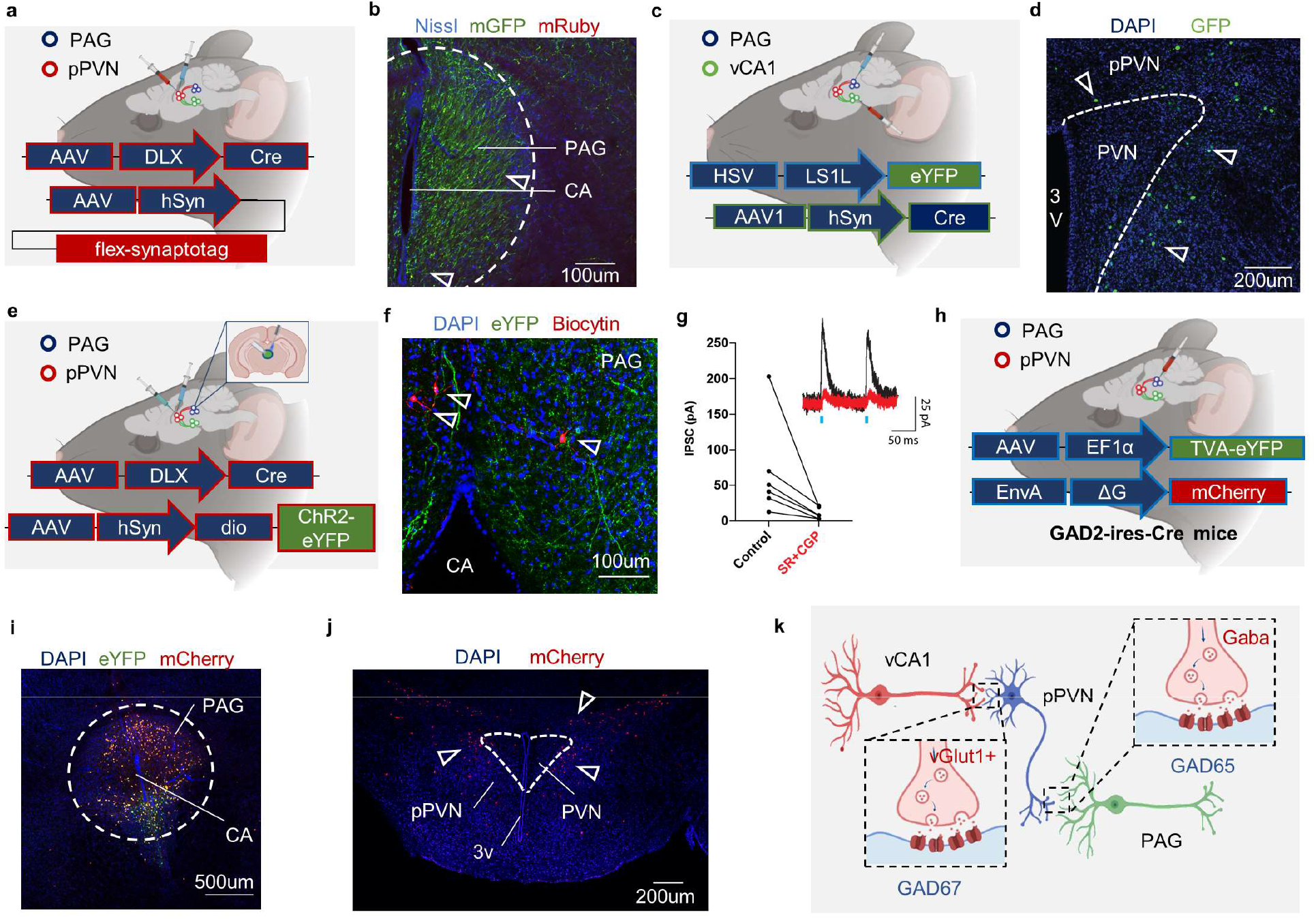
pPVN Halo Cells provide inhibition of GAD65^+^ inhibitory neurons of the PAG. **(a)** schematic of DLX driven synaptotag viral approach in the pPVN. **(b)** pPVN fibers (green) and terminals (red/yellow) are present in both the dorsolateral (dl) and ventrolateral (vl) PAG. **(c)** schematic of trans-synaptic high-titer AAV1-Cre approach for labeling pPVN relay cells between vCA1 and the PAG. **(d)** vCA1 projections to the PAG are routed through the pPVN. **(e)** schematic of optogenetic viral approach for whole-cell patch clamp. **(f)** representative image of biocytin (red) filled DAPI^+^ cells (blue) within ChR2-eYFP expressing fibers (green) in the PAG. **(g)** Quantification of PAG IPSC amplitude before and after application of 1 µM SR95531 and 2 µM CGP55845 (black circles are individual cells). **(h)** schematic of pseudotyped rabies virus approach on *GAD2-IRES-Cre* mice. **(i)** expression of TVA (green) and ΔG (red) expression in the PAG. **(j)** retrograde expression of mCherry (red) in the pPVN. **(k)** Disinhibition model for vCA1 vGlut1^+^ glutamatergic excitation of GAD67^+^ GABAergic inhibitory cells controlling GAD65^+^ GABAergic PAG neurons.

To examine if pPVN neurons receiving vCA1 inputs projected to the dlPAG, we injected a high-titer AAV1 expressing Cre recombinase (AAV1-Cre) into vCA1 followed by a Cre-dependent HSV expressing GFP (HSV-LS1L-GFP) in the PAG (Fig. 4c). High-titer AAV1 Cre recombinase exhibits a transsynaptic jump into downstream neurons (*40*) while HSV is taken up by neuronal terminals and retrogradely transported back to the nucleus (*41*). We identified pPVN cells that received Cre from vCA1 neurons and successfully expressed the eYFP from the HSV (Fig. 4c-d), thus confirming that some pPVN cells receiving vCA1 inputs project to the PAG. We next determined how pPVN inhibitory neurons influenced the PAG. We injected an AAV expressing channelrhodopsin (AAV8-hSYN-DIO-ChR2-eYFP) in the pPVN. Three weeks later we prepared acute slices of the PAG and obtained whole-cell patch clamp recordings of dlPAG neurons (Fig. 4f). Light stimulation of ChR2^+^ fibers in the PAG induced strong IPSCs in PAG neurons which were abolished by application of GABAergic synaptic blockers, confirming that the pPVN effectively inhibits PAG cells through GABA release (Fig. 4fg).

Previous studies have shown that long-range projections from the central amygdala to the dlPAG preferentially target inhibitory or excitatory cells thus leading to titrated fear responses (*42*). We used a rabies viral strategy to determine the identity of the PAG cells receiving pPVN inputs. GAD2-Cre or vGlut2-Cre mice were injected with a Cre-dependent rabies helper virus expressing eYFP followed by injection of a pseudotyped G-deleted rabies virus expressing mCherry (Fig. 4h & S5a) and examined the PAG and pPVN (Fig. 4i-j). GAD2-cre (Fig. 4i-j), but not vGLUT2-cre (Fig. S5b-c), mice exhibited a retrograde expression pattern that recapitulated the DLX-based (Fig. S4b) and vCA1→pPVN synaptophysin-mRuby (Fig. 1j & 2c) expression patterns. These data show that pPVN neurons project to inhibitory GAD2^+^ neurons in the PAG. Taken together, our findings reveal that vCA1 neurons excite pPVN inhibitory neurons which in turn silence PAG inhibitory neurons (Fig. 4k).

Finally, we sought to examine whether silencing vCA1→pPVN axon terminals (1) produced a sustained pattern of neural activation (i.e., c-Fos) in the PAG and (2) if more temporally precise vCA1→pPVN terminal silencing (Fig. 5a) would recapitulate our chemogenetic vCA1→pPVN cell body silencing experiments (Fig. 2k & m). We found that timed optogenetic silencing during acquisition of learned fear produced heightened responsivity to the first foot-shock (Fig. 5b-c) – an observation consistent with our chemogenetic findings. Moreover, vCA1→pPVN terminal silencing during conditioning selectively elevated c-Fos expression in the vlPAG, but not dlPAG (Fig. 5d-e). Similarly, optogenetic silencing of vCA1→pPVN terminals enhanced freezing during the early parts of exposure to the innately fearful predator odor TMT (Fig. 5f-g). However, optogenetic vCA1→pPVN terminal silencing globally elevated c-Fos throughout both the dlPAG and vlPAG (Fig. 5h-i). These data suggest that vCA1→pPVN circuit silencing regulates neural activity differentially in the dorsolateral and ventrolateral subdivisions of the PAG to titrate defensive responses to learned and innate threats (Fig. 5j).

**Figure 5.**
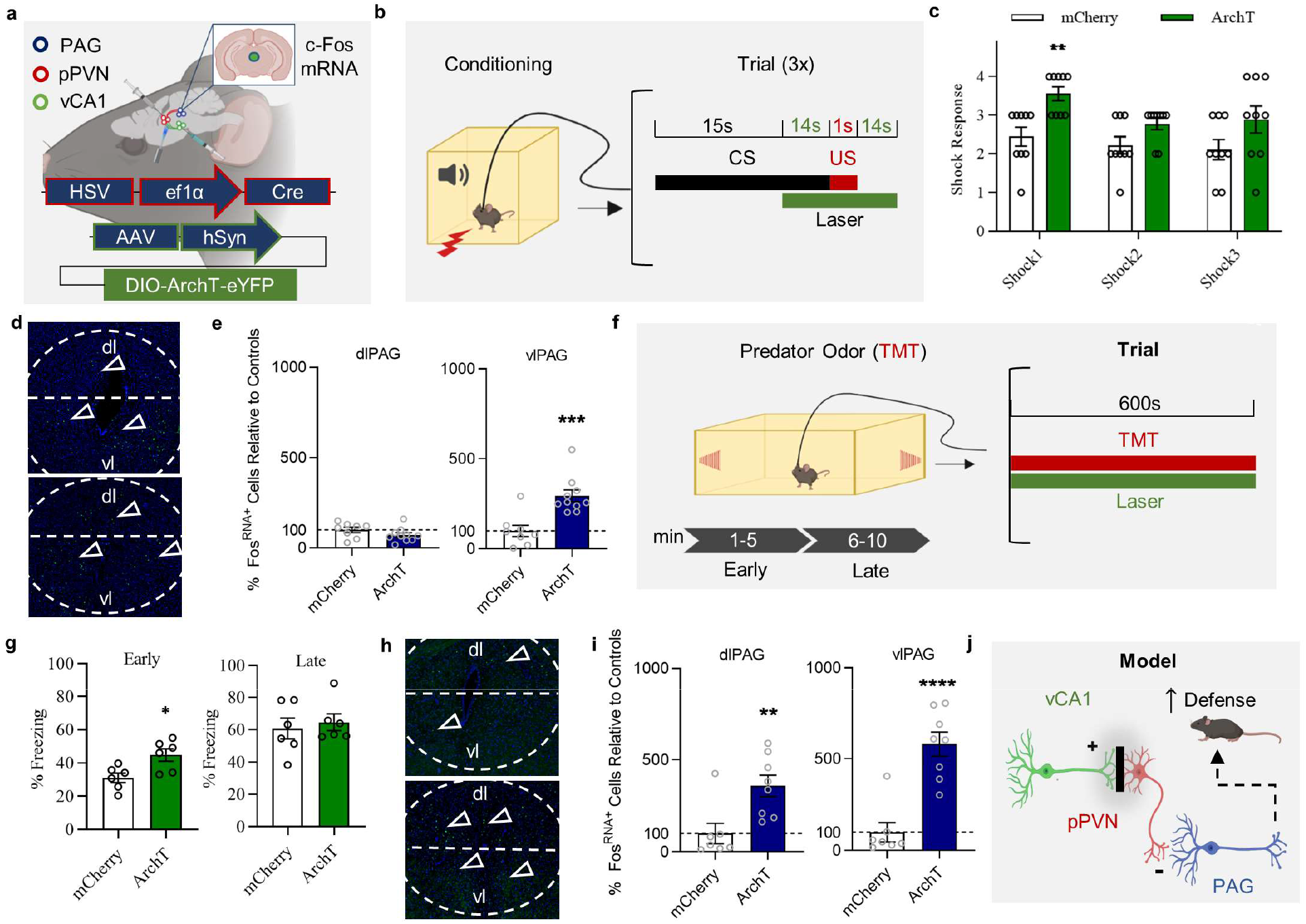
Optogenetic silencing of vCA1→pPVN terminals in the pPVN enhances responding to the first moments of innate threat and regulates neural activity in the PAG. **(a)** schematic of dual HSV and AAV optogenetic approach. **(b)** schematic of fear conditioning and optical stimulation parameters. **(c)** silencing during conditioning significantly suppressed shock responsivity to the first shock (Mann-Whitney_2-tailed_ U = 10, p = 0.0052), but not at shock2 (Mann-Whitney_2-tailed_ U = 21.5, p = 0.1189) and marginally at shock 3 (Mann-Whitney_2-tailed_ U = 19.5, p = 0.0693). **(d)** representative images of c-Fos expression in the PAG 1-hr after conditioning. **(e)** quantification of c-Fos mRNA in the dorsolateral PAG (dlPAG; left panel) and ventrolateral PAG (vlPAG) right panel. Silencing selectively elevated c-Fos in the vlPAG (t_16_=4.229, p=0.0006), but not dlPAG (t_16_=1.482, p= 0.1577). **(f)** schematic of innate predator odor TMT and optical stimulation parameters. **(g)** freezing during early and later parts of exposure to TMT. Silencing selectively enhanced freezing during the early (t_10_=2.902, p=0.0158), but not later parts of conditioning (t_10_=0.4680, p=0.6498). **(h)** representative images of c-Fos expression 1-hr following conditioning in the PAG. **(i)** quantification of c-Fos mRNA in the dlPAG (left panel) and vlPAG (right panel). Silencing elevated c-Fos in the dlPAG (t_13_=3.194, p=0.0070) and vlPAG (t_13_=5.656, p<0.0001). **(j)** Behavioral model for how silencing vCA1→pPVN terminals controls the PAG and defensive behavior. Bars and error bars are mean ± S.E.M, AFC opto fos: n_mCherry_=8 n_ArchT_=10, TMT: n_mCherry_=7 n_ArchT_=8, *p < 0.05, **p < 0.01, ***p < 0.001.

## Discussion

An enduring focus in the field of neuroscience is deciphering precisely how the ventral hippocampus controls learned fear, innate fear, and anxiety-like behaviors (*11, 19*). Using a combination of viral, electrophysiological, chemogenetic, fiber-photometric, and optogenetic approaches, we discovered a novel ventral hippocampal CA1 → hypothalamic peri-paraventricular circuit (vCA1→pPVN) which gates learned and innate fear, but not milder anxiety-like behaviors. This circuit operates via vCA1^vGlut1^ excitation of pPVN^GAD67^ inhibitory neurons. vCA1→pPVN circuit silencing of cell bodies (chemogenetics) or terminals (optogenetics) enhances active (foot-shock reactivity at conditioning) and passive (freezing to TMT) defensive behaviors. Moreover, *in vivo* fiber photometry revealed that the vCA1→pPVN circuit increases its activity during the first moments of a threat. Finally, we found that pPVN^GAD67^ neurons receiving vCA1 inputs routes information to dlPAG inhibitory neurons to provide long-range inhibition of pPVN^GAD65^ neurons. Importantly, this circuit influences neural activity in the PAG differently for learned vs. innate threats.

Recent neuroanatomical tracing data in mice has shown that ventral hippocampal CA1 neurons target multiple hypothalamic areas, including the peri-paraventricular nuclear area of the hypothalamus (*29*). Unlike its dorsal counterpart, vCA1 projects to a variety of limbic and cortical structures, including the medial prefrontal cortex (mPFC), basolateral amygdala (BLA), lateral hypothalamus (LH), and nucleus accumbens (NAcc; (*12, 13, 24, 43*). Our tracing data revealed little overlap between the vCA1→pPVN pathway and the vCA1→BLA, vCA1→mPFC, vCA1→NAcc, or vCA1→LH pathways (Fig. S1), suggesting that vCA1→pPVN neurons represent a relatively distinct vCA1 population.

Ventral hippocampal projections, including those to the lateral hypothalamus which control innate anxiety-like behaviors (but not learned contextual fear), are functionally segregated (*22*). For example, projections to the mPFC control anxiety-like behaviors whereas projections to the BLA control contextual fear (*10, 13*). However, even within the population of vCA1→BLA neurons, only a proportion selectively responds to the foot-shock (i.e., an unconditioned/innate stimulus). These neurons are triggered by the US irrespective of whether the CS is paired with a tone or a context (*44*). In fact, earlier work has shown that ibotenic acid lesions of the ventral hippocampus disrupt fear to foot-shock paired contexts and predator odor USs (*19*). However, previous studies which focused solely on short-lasting foot-shock USs likely neglect USs arising from other sensory modalities. Our study provides new insights into how the ventral hippocampus and discrete vCA1→pPVN projections control the magnitude of a defensive response to USs arising from auditory, olfactory, and tactile sensory modalities. Given that we did not detect any effects of chemogenetic silencing on measures of anxiety-like behavior (i.e., open field and elevated plus maze tests), sensory motor gating (outside of the highest startle pre-pulse), or short-term habituation (Fig. S3), our findings suggest that this vCA1→pPVN circuit dampens heightened defensive responses under conditions of elevated threat intensity. Therefore, it is possible that this circuit serves as an evolutionary adaptive gating mechanism for regulating diverse cortical and limbic inputs onto the PAG prior to executing an adaptive defensive response.

Recent computational models have sought to decipher how the hippocampus processes multimodal sensory information (*33*). A lingering question in the field is with how sensory information is sampled and processed across time when entering a new or familiar environment. Recent views, which assume serial and random stimulus sampling, suggest that the hippocampus may be probing sensory cues within the environment to select the correct configural or conjunctive representation (*33, 34*). In fact, our calcium recordings suggest that vCA1→pPVN projections show temporal activity patterns which match what the hippocampus is predicted to during the first moments of an experience (*33*). Moreover, along with short-duration calcium activity during tactile foot-shocks and sustained calcium activity during long-duration exposure to an innate olfactory threat, our data suggest vCA1→pPVNs are at least partly involved in multimodal sensory processing and, importantly, these cells parcel out sensory stimuli depending on factors related to threat intensity (low-level anxiety vs. intense fear), threat duration (scaling on the order of seconds to minutes), and threat type (learned vs. innate).

Neurons in the pPVN (i.e., Halo Cells) are primarily GAD67^+^ inhibitory neurons. Halo Cells send long-range projections to PAG^GAD65^ inhibitory neurons. Neuronal excitation of inhibitory pPVN^GAD67^ neurons by vCA1^vGLUT1^ neurons likely increase inhibition of PAG^GAD65^ inhibitory neurons– presumably increasing excitatory transmission (Fig. 4) and neural activity in the PAG (Fig. 5) as well as enhanced defensive responding (Fig. 2 vs 5). Past work indicates that dorsolateral divisions of the PAG control active avoidance whereas the ventrolateral division control passive defensive (e.g., freezing; (*45*)). Recent work has shown that defensive behavior is tightly regulated via disinhibition of the PAG by the central amygdala (*42*). That is, GABAergic central amygdala projections to PAG^GAD65^ neurons locally disinhibit PAG^vGlut2^ neurons. These PAG^vGlut2^ neurons then directly project to pre-motor neurons to influence defensive responding to threats (*42*). Thus, the vCA1^vGlut1^→pPVN^GAD67^→PAG^GAD65^ circuit likely functions to disinhibit the PAG^vGlut2^ when threat-intensity is high (Fig. 2 vs. Fig. S3). More work is needed to understand how amygdala and hippocampal (*46*) networks jointly coordinate defensive responses to produce maladaptive behaviors.

Accumulating evidence suggests that (1) the amygdala and hippocampus are key sites in many anxiety and stress-related disorders (*47*) and (2) pre-existing hippocampal changes (evidenced by smaller volume) may underlie the emergence of disorders like post-traumatic stress disorder (PTSD). In fact, a causal factor in the development of PTSD is heightened defensive responses (*48*). Several studies in recent years have sought to identify the neural circuit mechanisms of anxiety and post-traumatic stress disorders. Our study provides some of the first mechanistic insights into how coordinated regulation of PAG activity by the ventral hippocampus may titrate how the initial moments of a threat are experienced and behaviorally represented. Future studies are needed to understand how amygdala and Halo cell circuits may work together to regulate how threats are initially processed, stored, and retrieved – as this may provide critical insights into how maladaptive behaviors emerge in disorders like post-traumatic stress disorder (PTSD).

## Acknowledgments

We thank Larry Swanson and Jeffrey B. Rosen for helpful experimental and theoretical insights. We thank Kevin Karl for help in building the elevated plus maze. We thank Amsal Madhani and Anna Rekow for help with the initial immunohistochemical experiments. We greatly thank Christina Doyle for help in acquiring viral constructs, laboratory reagents, and overall support.

## Author Contributions

A.A. and E.R.K conceived the study. A.A. performed chemogenetic experiments. A.A. performed optogenetic experiments. A.A. performed in vivo fiber photometric calcium recordings. A.A. performed in situ hybridization. A.A. with R.N. re-designed the AAV-DLX construct. A.A. and F.L. performed whole-cell patch clamp experiments. A.A., C.P., and F.L. performed surgeries. A.A., M.N.F. and F.L. performed the immunohistochemistry. R.N. produced all HSV constructs. J.B.R., C.A.D, A.K. and L.K.F. provided experimental insight. A.A., F.L. and E.R.K analyzed data. A.A. and E.R.K. wrote the manuscript.

## Data and Materials Availability

The AAV-DLX-Cre-eYFP construct is available from A.A. and/or R.N. All other data are available upon reasonable request.

## Materials and Methods

### Subjects

10–18-week-old C57/BL6, GAD2-Cre, or vGlut2-Cre mice were used for all experiments (Jackson Laboratory, Bar Harbor, ME). Same sex mice were housed 4 to a cage on a 12h light/dark cycle (lights on from 7:00 – 19:00) with ad libitum access to food and water. Mice were acclimated to the colony for at least 1-week prior to the start of experimentation. All experimental sessions were conducted during the light phase between 09:00 and 17:00 h. Procedures were conducted in accordance with the US National Institute of Health Guide for the Care and Use of Experimental Animals and were approved by the Columbia University Institute of Comparative Medicine and the New York State Psychiatric Institute Department of Comparative Medicine.

### General Surgery Procedure

For all surgeries, mice were weighed and then anesthetized by either isoflurane gas or a cocktail of ketamine/xylazine (100 mg/kg and 10 mg/kg respectively). Local (e.g., marcaine) and general analgesics (e.g., carprofen) in addition to eye lubricant was administered to minimize any pain or distress during surgery. Following anesthesia, the animal’s head was mounted into a stereotax and the surgical site was triple cleaned with 70% ethanol and betadine prior to any incision. A craniotomy was performed to remove the skull directly above the target site. The instrument (e.g., glass-pulled pipette, fiber optic cable, ferrule, etc.) was then lowered to the target coordinates within the brain. All viral injections were performed with either a NanoJect II or III nanoliter injector (Drummond Scientific, Bromall, PA) at 23 nL/s or 4 nL/s, respectively. After the procedure, mice were sutured and triple antibiotic was applied to the surgical site and the mouse was housed in a clean cage with its original cage mates (who have already undergone surgery).

### Viral Injections

Mice received either unilateral or bilateral, single or double viral injections. Injections were delivered at the following coordinates relative to Bregma (in mm): ventral hippocampal CA1 subfield (vCA1): AP = −2.9, ML = ±2.4, DV = −4.6, the peri-paraventricular nucleus of the hypothalamus (pPVN): AP = −1.06, ML = ±0.2, DV = −5.35, or the periaqueductal grey (PAG): AP = −4.4, ML = ± 0.5, DV = −2. Unless otherwise specified, 200 nL of virus was injected into the target region. The following viruses were used: bilateral AAV5-CAMKIIα −eYFP in vCA1, unilateral HSV-EF1α-GCaMP6f(HT) in pPVN, HSV-EF1α-EYFP-IRES-Cre(HT) in pPVN, HSV-EF1α-Cre(HT) in pPVN, or HSV-EF1α-mCherry-IRES-Cre in pPVN (HT), AAV2/9-DLX-Cre-eYFP (Rachael Neve, Viral Gene Transfer Core, MGH). For experiments involving double injections, two weeks after the first injection, mice also received AAV-hSyn-flex-mGFP-2A-Synaptophysin-mRuby in vCA1 (Synaptotag: Stanford Vector Core), AAV2/8-Syn-dio-mCherry in vCA1 (control, Addgene), hM4Di (AAV2/8-Syn-DIO-hM4DI-mCherry; Addgene), AAV2/9-EF1α-dio-ChR2(E123A)-EYFP-WPRE-hGH (channelrhodopsin, Penn Vector Core), AAV2/9-EF1α-dio-EYFP-WPRE-hGH (control, Penn Vector Core). For rabies injections, mice received 50 nL of helper AAV2/8 syn.dio.TVA.2A.GFP.2A.B19G (UNC vector core) followed 2 weeks later by 300nl of rabies SAD.B19.EnvA.ΔG.mCherry (SAD-B19 strain, Addgene Cat# 32636 prepared by the Salk institute vector core) in the PAG.

### Fiber Optic Implants

For fiber photometry experiments, a single 400 µm optic ferrule was implanted approximately 200 µm above vCA1. For optogenetic experiments, custom 200 μm fiber optic ferrules (Precision Fiber, MM-FER2007C-2300-P) were bilaterally implanted above axon terminals in the pPVN.

For optogenetic experiments, 200 nL of HSV-hEF1α −Cre (MIT, RN425) was injected bilaterally into the pPVN and allowed to express for two to three weeks before 200 nL of AAV5-CAG-FLEX-ArchT-tdTomato (Addgene, 28305) (or pAAV2-hSyn-DIO-mCherry (Addgene, 50459) was injected into vCA1. Two weeks later, optical fibers were bilaterally implanted above the pPVN (AP −1.06, ML ±2.00, DV - 5.35, angle ±18°). This viral strategy allowed us to selectively and transiently silence the terminals of neurons projecting directly from the hippocampus (vCA1) to the pPVN using light stimulation.

Optical fibers were constructed in-house using 200 μm optical fibers (ThorLabs, FT200UMT) and 230 μm ferrules (Precision Fiber Products Inc., MM-FER2007C-2300-P). Before implantation, the power output of each fiber was tested to eliminate fibers with coupled outputs outside of the acceptable range and pair fibers for implantation into animals. During implantation surgery, three additional holes drilled into the skull surface were used for anchoring fibers to screws. Once the fibers reached the target location, super glue and accelerant were applied to the area to anchor the fiber/ferrule to the stabilizing screws. The entire area was then covered in glass ionomer (GC, FujiCEM 2).

### Slice Preparation

Mice were killed under isoflurane anesthesia by perfusion into the right ventricle of an ice-cold solution containing the following (in mM): 10 NaCl, 195 sucrose, 2.5 KCl, 10 glucose, 25 NaHCO_3_, 1.25 NaH_2_PO_4_, 7 Na-pyruvate, 0.5 CaCl_2_, and 7 MgCl_2_. The skull was placed in the same ice-cold medium and the brain was removed carefully from the skull. The brain was then glued upright with the dorsal side facing the blade and a small block of 4% agar was placed against the ventral side for mechanical stabilization. 400 µm thick coronal slices were prepared with a vibratome (VT1200S, Leica) in the same ice-cold dissection solution. Brain slices were then transferred to a chamber containing 50% dissecting solution and 50% ACSF (in mM: 125 NaCl, 2.5 KCl, 22.5 glucose, 25 NaHCO_3_, 1.25 NaH_2_PO_4_, 3 Na-pyruvate, 1 ascorbic acid, 2 CaCl_2_ and 1 MgCl_2_). The chamber was kept at 34°C for 30 min and then at room temperature for at least 1h before recording. Dissecting and recording solutions were both saturated with 95% O2 and 5% CO2, pH 7.4.

### Electrophysiological Recordings

Slices were mounted in the recording chamber under a microscope. Recordings were acquired using a Multiclamp 700 A amplifier (Molecular Device), data acquisition interface ITC-18 (Instrutech) and the Axograph X software. Whole-cell current-clamp recordings were obtained from LS cells with a patch pipette (4–5 MΩ) containing the following (in mM): 135 K methylsulfate, 5 KCl, 0.2 EGTA-Na, 10 HEPES, 2 NaCl, 5 ATP, 0.4 GTP, 10 phosphocreatine, and 5 μM biocytin, pH 7.2 (280–290 mOsm). The liquid junction potential was 1.2 mV and was not corrected. Voltage-clamp recordings were performed with an intracellular solution containing 135 Cs methylsulfate instead of K methylsulfate. Series resistance (15–25 MΩ) was monitored throughout each experiment; cells with a >20% change in series resistance were discarded. For light stimulation, pulses of blue light (pE-100, Cool LED) were delivered through a 40× immersion objective and illuminated an area of 0.2 mm^2^. The illumination field was centered over the recorded cell.

### Drugs

For chemogenetic experiments, clozapine-N-oxide dihydrochloride (CNO; Tocris Biosciences) was dissolved in saline to 10 mg/kg. Prior to behavioral experiments, the solution was diluted to 5mg/kg and administered to mice via intra-peritoneal injections 20 minutes prior to the start of behavioral sessions. For whole-cell patch clamp experiments, we used the following drugs from Tocris: in a subset of experiments the following drugs were used at the following concentrations via bath application: SR95531 (1 μM, #1262), CGP55845 (2 μM, #1248), D-APV (50 μM, #0106), CNQX (20 μM, #1045). Drugs were bath applied following dilution in ACSF.

### In Vivo Fiber photometry

Animals were implanted with custom 400 μm optical ferrules above vCA1 (−2.9 Bregma; Doric Lenses, Quebec, CA). Light was delivered at a final intensity of 2.24 mW (473 nm) and 2.76 mW (405 nm) at the tip of the patch-cord before an ∼60% reduction in final power output prior to coupling with the implanted ferrule. Prior to behavioral testing, mice were habituated to the experimenter and coupler for two days.

Fiber photometry experiments were conducted similar to our previous work (*14*). A 405 nm and 463 nm LED were coupled to a fluorescence mini-cube (FMC). The sample port was connected to a 1 × 1 fiber optic rotary joint to deliver light into optical fibers permanently implanted above ventral CA1 during behavior. Light between 420-450 nm or 500-540 nm were collected through the FMC on separate Newport 2151 photo-receiver modules. The fluorescent signals were collected in AC-high mode and converted to voltage via the formula V = PRG, where V is collected voltage, P is the optical input power in watts, R is photodetector responsivity in amps/watts (0.2 – 0.4), and G is the trans-impedance gain of the amplifier. Raw signals for 463 nm excitation (GCaMP6f) and 405 nm excitation (background auto-fluorescence) were recorded and processed using Doric Neuroscience Studio software. Raw signals were low-pass filtered and ΔF was calculated via a time fitted running average for the 473 nm (F_1_) and separately the 405 nm (F_2_) channel. ΔF/F = F_1_ −F_2_ / F_2_ and data for each animal were further transformed by employing a peak enveloping Fourier transform and z-score normalization before collapsing animals of a group together. All analysis was conducted on z-score normalized data using Matlab (Mathworks).

### Optogenetic Behavioral Manipulations

For optogenetic studies, 532nm green laser light was delivered continuously (Laserglow Technologies). Light was delivered during a 30-s block surrounding the shock at fear acquisition during delay auditory fear conditioning or for the entire 600-s session during predator odor exposure. Following optogenetic experiments, animals were sacrificed 1-h later via rapid decapitation for RNAscope experiments to measure c-Fos mRNA.

### General Behavioral Procedure

For all behavioral procedures, mice were transported in their home cage to the testing room for 30-m prior to any testing or drug injections. Different behavioral tasks were run in different rooms and groups were counterbalanced within and across days to control for any experimenter or test-order effects.

### Auditory Fear Conditioning

Fear conditioning occurred over a 3-day period using a Med Associates .4-chamber NIR video fear conditioning system. On day 1, mice were exposed to a conditioning chamber (Med Associates) for a 120-s baseline period (baseline) followed by three 30-s 5000 Hz tones that co-terminated with a 1-s 0.4mA foot-shock. Chambers Following the last tone + foot-shock pairing, mice remained in the chamber for an additional 60-s. On day 2, mice were re-exposed to the conditioning chamber for 300-s. On day 3, mice were introduced into a novel context that differed from the conditioning context in terms of lighting, spatial dimensions, olfactory components, and tactile cues. Following a 120-s baseline period, mice were exposed to five 30-s 5000 Hz tones. Freezing was measured via integrated Med Associates Video Freeze software with an episode of freezing defined as immobility for 75% of 30 frames in a 1-s time bin.

### Shock Responsivity

Shock responsivity was scored using procedures outlined by (*49*) similar to our previous work (*31*). Briefly, responses were videotaped and scored offline by an experimenter on an ordinal scale from 0-4 (0 = no response, 1 = flinch, 2 = hop, 3 = horizontal jump, and 4 = vertical jump).

### Open Field (OF) Behavior

Open field behavior was run on the SmartFrame open field system from Kinder Scientific (Poway, CA). All mice were placed in the center of the open field arena (50 cm × 50 cm × 50 cm) and allowed to freely explore the arena for a 5-min period. The center was defined as a square area approximately 12.5 cm from each wall. Motor monitor software was used to examine the time spent in the periphery, time spent in the center, and the total distance traveled in each area. The chamber was cleaned with 70% ethanol between tests.

### Elevated Plus Maze (EPM) Behavior

Elevated plus maze behavior was conducted in a maze comprised of a 5 cm × 5 cm center, two closed 50 cm × 10 cm × 40 cm arms, and two open 50 cm × 10 cm arms. The maze was elevated 50 cm above the ground. Recorded videos were analyzed using Anymaze behavioral software in order to score time spent in the closed arms, time spent in the open arms, time spent in the center, and distance traveled.

### Pre-pulse Inhibition

Pre-pulse inhibition was conducted via the SR-Lab Startle Response System from San Diego Instruments using a protocol identical to previous work (*50*). Each session included a 65-dB background noise throughout the session and started with a 5-m acclimation period. All stimuli were presented with an ITI ranging from 7 s to 23 s with a 15 s average. The max startle amplitude over the 40-ms recording window was used for analyses. Following acclimation, mice were presented with five blocks of acoustic startle stimuli. In block one, mice received five 40-ms 120 dB startle pulses. In block two, mice received four 40-ms pseudorandomized presentations of each 80 dB, 90 dB, 100 dB, 110 dB, and 120 dB. In block three, mice were tested with 12 PPI presentations of 68 dB, 71 dB, and 77 dB pre-pulses that preceded that 120 dB stimulus in addition to interspersed non-PPI 120-dB pulse alone trials. In block 4, mice were tested identically to block one.

Short-term startle habituation was calculated by the formula: %Δ Habituation = ((Block 1 – Block 2) / Block 2) × 100. Percent change in PPI was calculated using the formula: % Δ PPI = ((100 - PPI Startle Response / 120 dB non pre-pulse trial)*100).

### Predator Odor Exposure

For odor exposure experiments, mice were exposed to 150 μmoles of the predator odor 2,5-dihydro-2,4,5-trimethylthiazoline (TMT) or 150 μmoles of acetic acid as a control odor in a similar manner to our previous work, see (*51, 52*)). Briefly, odor was added to pieces of triangular filter paper on opposite sides of a distinct rectangular chamber (∼ 30 cm × 24 cm × 9 cm) positioned inside of a fume hood immediately prior to placement of the mouse. Upon placement, mice freely explored the chamber for 10-m. At the end of the session, mice were placed into a new chamber with tested littermates and housed in a distinct room for 24-h to allow for any residual odor to clear prior to returning mice to the colony.

### Immunohistochemistry

For perfusions, mice were first anesthetized with a Ketamine/Xylazine cocktail (200mg/kg and 20mg/kg respectively) prior to transcardial perfusion with 10 mL of ice-cold 1x PBS (7.4 pH) followed by 10 mL of 4% PFA. Brains were removed and post-fixed in 4% PFA for 18h. Following post-fixation, brains were subjected to two 30-m washes in 1x PBS, prior to slicing. Brains were then embedded in a 4% agarose/1x PBS w/v solution prior to mounting and slicing on a vibratome (Leica VT 1000 S). 40 μm or 60 μm sections containing the pPVN (−0.70 to −1.06 mm relative to Bregma), vCA1 (−2.80 to −3.28), or PAG (−3.16 to −3.80) regions.

For staining, sections were washed twice for 10-m in 1x PBS and then incubated for 1-hr in a blocking buffer containing 3-5% Normal Goat Serum (Jackson Immuno Research, 005-000-121) and 0.01-0.03% Triton-X (Sigma, X100), made up in 1x PBS. Sections were co-labeled for GFP & RFP using anti-chicken GFP (Aves, GFP-1020) and anti-rabbit RFP (Abcam, 167453), each at a concentration of 1:1000, diluted in blocking buffer for a 48h incubation at 4°C.

Sections were then placed in three 10-m washes in 1x PBS. Brain tissue was stained with secondary antibodies complementary to the primaries used; goat anti-chicken Alexa Flour Plus 488 IgY (Invitrogen, a32931) and goat anti-rabbit Alexa Flour 633 IgG (Invitrogen, a21071), each at concentrations of 1:1000 in blocking buffer. DAPI was used for nuclear staining at a concentration of 1:10,000. Sections were placed in two 30-m washed in 1x PBS before being replaced in fresh 1x PBS for mounting. Sections were mounted onto Superfrost Plus Microscope Slides (Fisher Scientific, 12-550-15), briefly allowed to dry, and sealed using Prolong Diamond Antifade Reagent (Thermo Fisher, P36961). No Primary controls were also used alongside optimization of the staining procedure. Slides were cured in the dark for 24-hr before being moved to −20°C for long-term storage.

### In Situ Hybridization

In situ hybridization was conducted using the RNAscope assay from ACDbio similar to our previous work (*31*). 16-20 μm sections were prepared on a cryostat and stored at −80°C prior to staining. For staining, slices were removed from the −80°C and immediately placed in a 10% neutral-buffered formalin solution for 15-m prior to subsequent dehydration in 50%, 70%, and 100% ethanol (twice) diluted in DEPC-treated water. A hydrophobic barrier was drawn around tissue sections following the last ethanol rinse and sections were incubated in Protease IV for 25-m prior to rinsing in DEPC-treated PBS. Sections were then incubated with their respective probes for 2-hr at 40°C. Followed by subsequent hybridization steps at 40°C with AMP1-4FL for 30-m, AMP2-FL for 15-m, AMP3-FL for 30-m, and AMP4-FL for 15-m. Sections were washed twice with RNAscope wash buffer following manufacturer’s instructions between hybridization steps. Following the last wash, slices were briefly counterstained with DAPI and prolong-gold antifade reagent (Thermo Fisher, P36961) was added prior to coverslipping. Negative controls were also run along with assays. Multiplex RNAscope for the PAG was conducted using probes targeting: c-Fos (mm-Fos-C2, 316921-C2), vGlut2 (mm-Slc17a6-C3, 319171), and GAD2 (mm-Gad2-C1, 439371) were used. For vCA1 and pPVN, probes targeting: GAD1 (mm-Gad1-C3, 400951), EGFP (mm-egfp-C2, 400281), vGlut1 (mm-Slc17a7-C1, 416631), or CRH (mm-Crh-C2, 316091) were used. All sections were counterstained with DAPI prior to cover-slipping and imaged within 2-days of the assay.

### Imaging

All images were captured on an Olympus FV1000 confocal microscope, Leica LSM 700 confocal microscope, a Nikon AZ100 Axioscan Multizoom slide scanner, or a Keyence BZ-X800. For co-localization experiments, tiled images were first captured on the Keyence BZ-X800. Any sections that exhibited overlap between fluorescent signal were then captured on an Olympus FV1000 to confirm co-localization of the signal. Images were captured at varying magnifications including: 10x, 20x, and 40x. Cell counts for immunohistochemical co-localization experiments were obtained via the ImageJ Cell Counter Plugin. For axon terminal analysis experiments and RNAscope studies, images were cropped and masked using Cell Profiler 3.0 to identify terminals or RNA within the PVN or pPVN or PAG (*53*). Schematics were created with Biorender.

### Statistics

Homogeneity of variance was first tested between groups and tests were corrected for any violations. For c-Fos counts and two-group behavioral analyses, independent samples t-tests were used to compare differences. For shock responsivity testing, Mann-Whitney U tests were used. For pre-pulse inhibition, a two-way ANOVA was used. For calcium recordings, one-way or two-way ANOVAs were used to analyze main effects on mean z-score transformed ΔF/F.

## Supplementary Figure 1

**Supplementary Figure 1.**
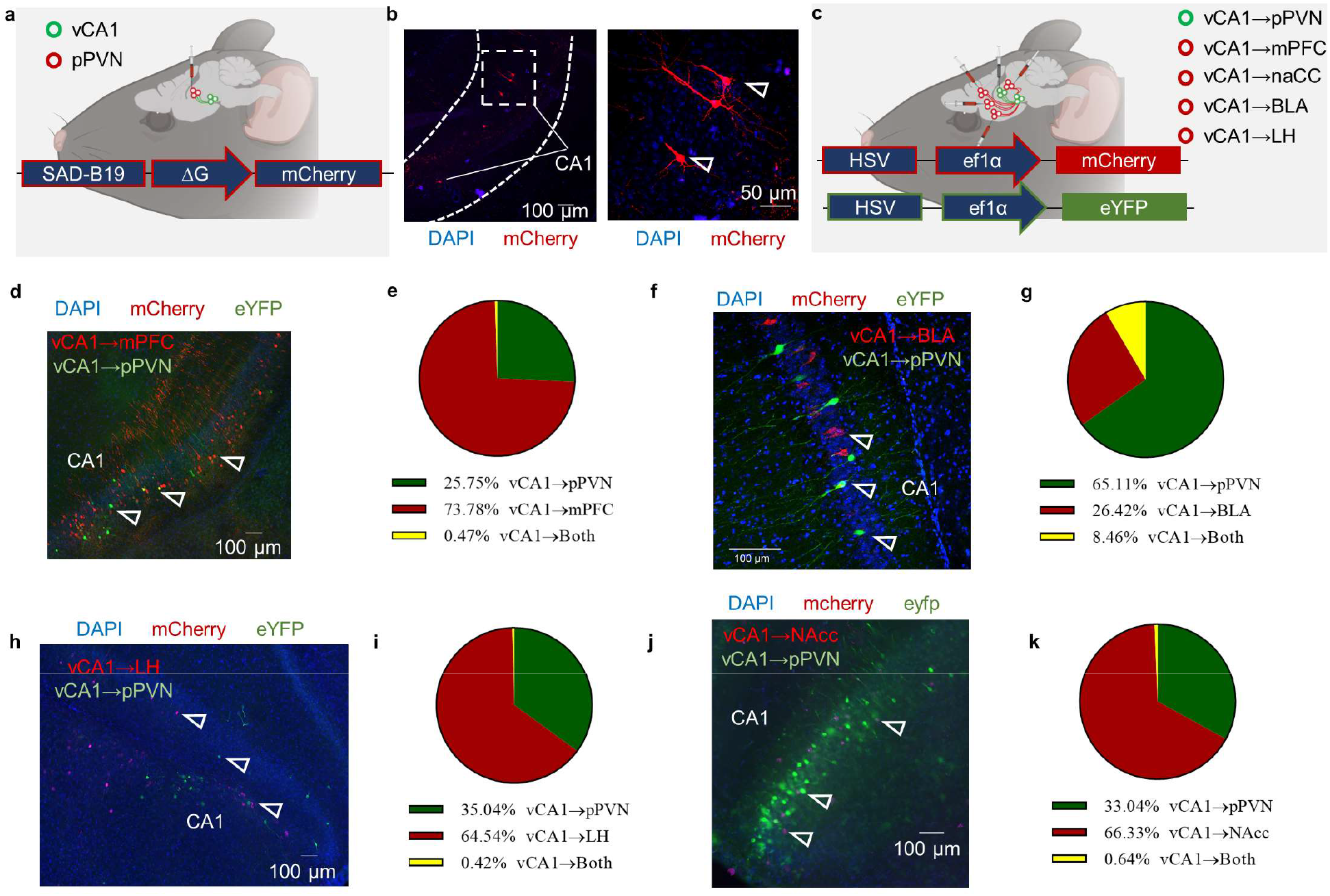
vCA1→pPVN projections are unique relative to other known projections of the ventral hippocampus. **(a)** schematic of SAD-B19-mCherry rabies virus injection into the pPVN. **(b)** right panel: representative image of mCherry^+^ rabies cells in the ventral hippocampus showing that projections to the pPVN area are monosyaptic, left panel: magnified image of mCherry^+^ cells in vCA1. **(c)** schematic of unilateral dual viral approach for circuit tracing. HSV-EF1α-mCherry was injected into known projection targets of the ventral hippocampus including: the medial prefrontal cortex (mPFC), nucleus accumbens (NAcc), basolateral amygdala (BLA), or lateral hypothalamus (LH). HSV-EF1α-eYFP was injected into the pPVN. Co-localization of eYFP and mCherry in the ventral hippocampus CA1 subfield was assessed following each injection. **(d)** representative ventral hippocampus image following injection of into the mPFC (mCherry) and pPVN (eYFP). **(e)** For mPFC (mCherry) and pPVN (eYFP) injections, 25.75% of labeled ventral hippocampal cells were eYFP^+^, 73.78% of labeled cells were mCherry^+^, and 0.47% of labeled cells expressed both. **(f)** representative ventral hippocampus image following injection into the BLA (mCherry) and pPVN (eYFP). **(g)** For BLA (mCherry) and pPVN (eYFP) injections, 26.42% were mCherry^+^, 65.11% were eYFP^+^, and 8.46% were both. **(h)** representative ventral hippocampal image following injection into the LH (mCherry) and pPVN (eYFP). **(i)** For LH (mCherry) and pPVN (eYFP) injections, 64.54% were mCherry^+^, 35.04% were eYFP^+^, and 0.42% were both. **(j)** representative ventral hippocampal image following injection into the NAcc (mCherry) and pPVN (eYFP). **(k)** For NAcc (mCherry) and pPVN (eYFP) injections, 66.33% of labeled ventral hippocampal cells were mCherry^+^, 33.04% of labeled cells were eYFP^+^, and 0.64% of labeled cells were both. n = 4 mice/experiment.

**Supplementary Figure 2.**
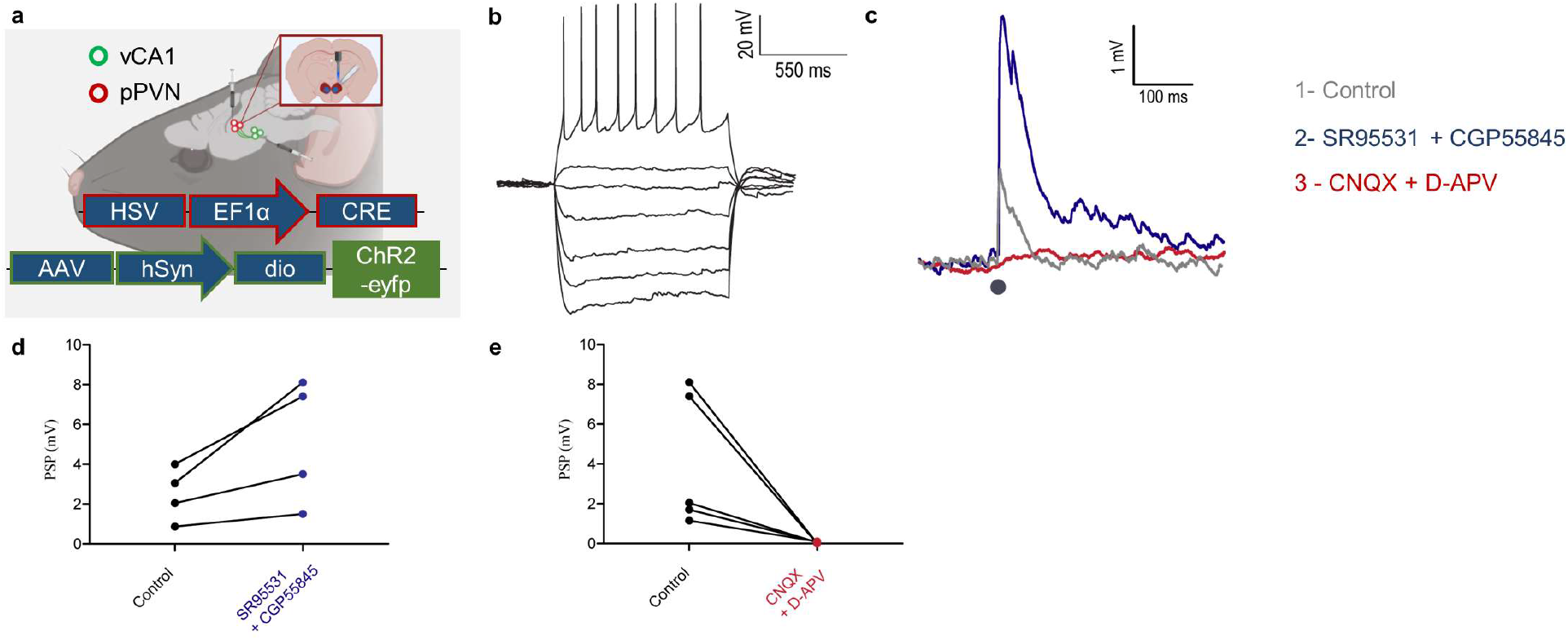
Optogenetic whole-cell patch clamp of pPVN neurons approach following injections of HSV-Cre-mCherry into the pPVN followed by AAV-DIO-ChR2-eYFP injection into vCA1. **(a)** Voltage response of a pPVN neuron during injection of current pulses. Magnification of inset from Fig. 1n (b) Representative voltage response of a pPVN neuron to light stimulation before and after application of 1 µM SR95531 and 2 µM CGP55845 then 20 µM CNQX and 50 µM D-APV. (c) Quantification of pPVN neurons post synaptic potential (PSP) amplitude before and after application of SR95531 andCGP 55845 to block GABAergic transmission (black circles are individual cells). (d) Quantification of pPVN post synaptic potential amplitude before and after application of CNQX and D-APV to block glutamatergic transmission (black circles are individual cells).

**Supplementary Figure 3.**
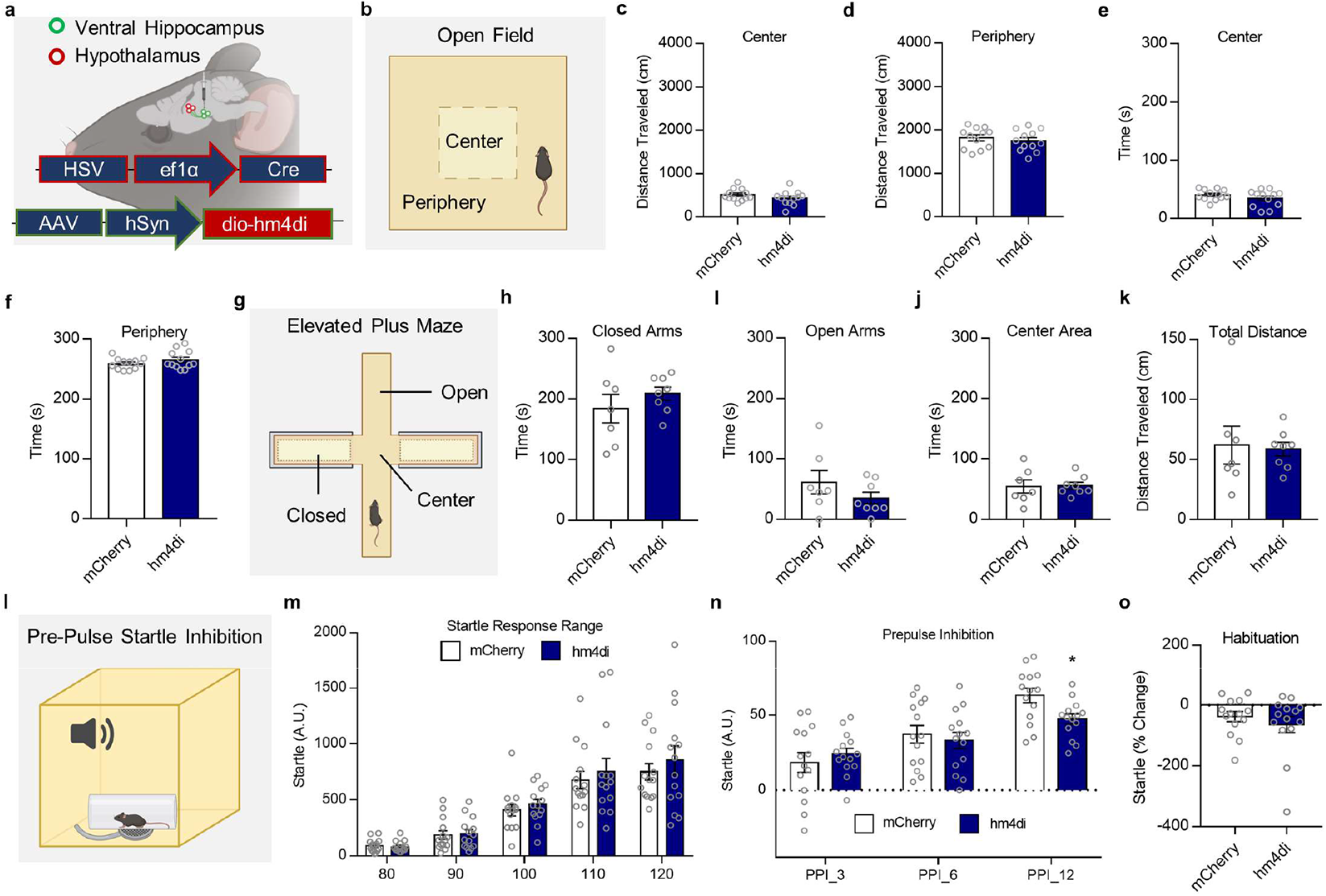
Chemogenetic silencing has no effect on open field (OF) or elevated plus maze (EPM) behavior, but selectively disrupts the highest intensity pre-pulse in pre-pulse inhibition (PPI). **(a)** truncated schematic representation of viral approach. **(b)** schematic of OF paradigm. **(c-d)** silencing has no effect on distanced traveled in the center (t_22_=1.234, p=0.2302) or periphery (t_22_=0.6727, p=0.5081) of the OF. **(e-f)** silencing has no effect on time spent in the center (t_22_=1.201, p=0.2424) or periphery (t_22_=1.237, p=0.2290) of the OF. **(g)** schematic of elevated plus maze. **(h-j)** silencing had no effect on time spent in the closed (t_13_=1.006, p=0.3326), open (t_13_=1.256, p=0.2313), or center areas (t_13_=0.1100, p=0.9141). **(k)** silencing had no effect on total distance traveled (t_13_=0.2184, p=0.8305) in the EPM. **(l)** schematic of PPI paradigm. **(m)** silencing had no effect on startle responsivity to tones alone (ANOVA_2-way Startle dB_: F_1.623,42.21_ = 57.15, p<0.0001). **(n)** silencing preferentially disrupted the highest intensity pre-pulse (ANOVA_2-way Time X Group interaction_: F_2,51_ = 19.61, p=0.0011). **(o)** silencing had no effect on short-term habituation (t_26_=0.8176, p=0.4210). Bar graph are mean ± S.E.M, Open Field: n_mCherry_=12 n_hm4di_=12, EPM: n_mCherry_=7 n_hm4di_=8, Startle: n_mCherry_=14 n_hm4di_=14, *p < 0.05.

**Supplementary Figure 4.**
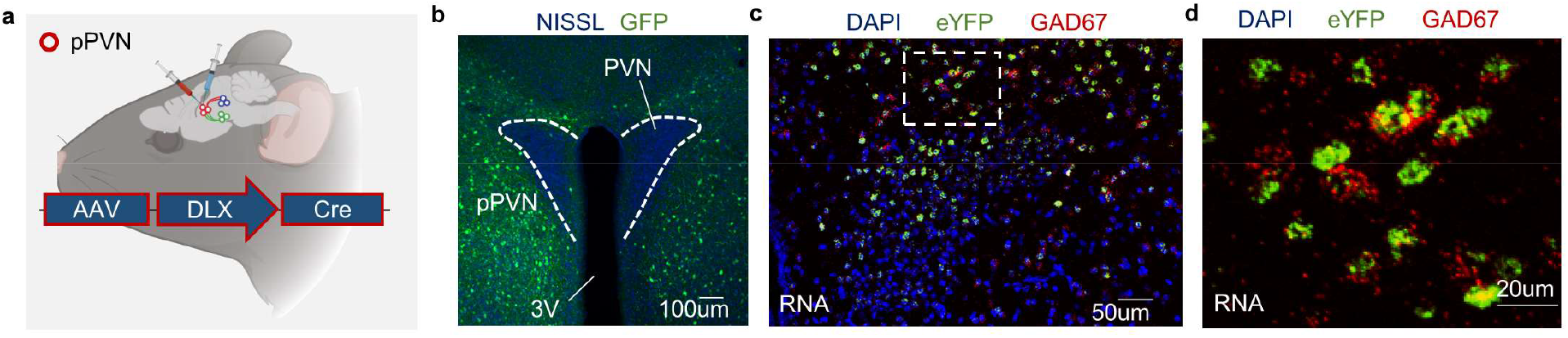
AAV-DLX-Cre-eYFP targets GABAergic pPVN neurons. **(a)** schematic of AAV-DLX-Cre construct injected into the pPVN. **(b)** AAV-DLX recapitulates the halo-like neuroanatomical pPVN pattern around the PVN. **(c)** In situ hybridization image showing the overlap between eYFP from the AAV-DLX-Cre-eYFP (green) and GAD67 (red). **(d)** magnified view of eYFP and GAD67 co-localization.

**Supplementary Figure 5.**
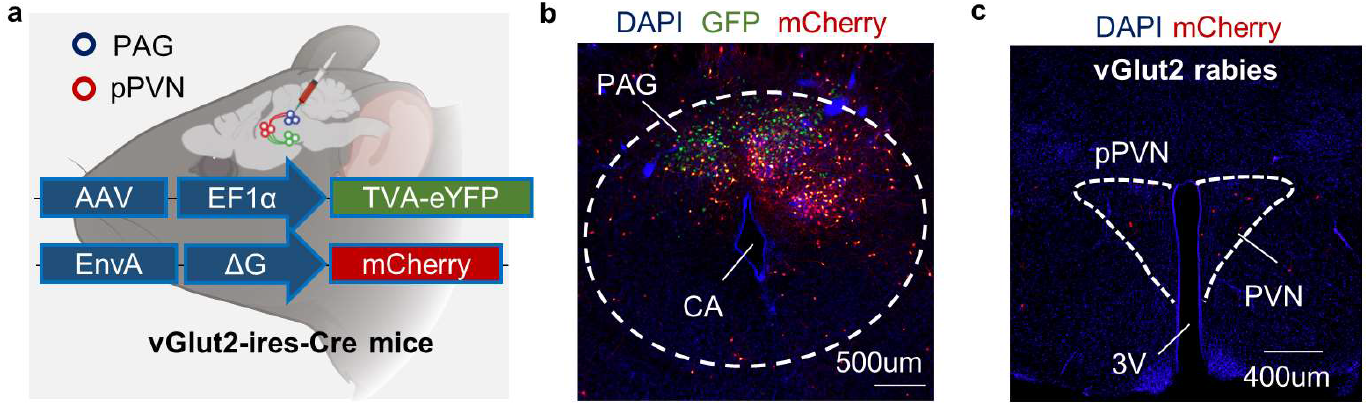
pPVN projections do not synapse onto PAG glutamatergic cells. **(a)** schematic for retrograde pseudotyped rabies tracing in *vGlut2-IRES-Cre* mice. (b) representative image of GFP and mCherry signal in PAG. (c) representative image of sparse rabies expression in the PVN, but not in pPVN.

